# Premovement suppression of corticospinal excitability is modulated by reaction time task requirements

**DOI:** 10.64898/2026.04.27.721107

**Authors:** Anthony N. Carlsen, Cassandra M. Santangelo, Christin M. Sadler, Dana Maslovat

## Abstract

The amplitude of motor-evoked potentials (MEPs) elicited using transcranial magnetic stimulation (TMS) has been shown to decrease in the short interval prior to response initiation. The cause of this premovement MEP suppression is currently unclear and has been attributed to various processes such as preparation-related inhibition preventing the premature release of planned action or increasing signal-to-noise ratio to facilitate rapid response initiation. The present study explored whether the decrease in MEP amplitude is affected by the task requirements, using reaction time (RT) paradigms that differ in the timeline of preparation and initiation of a motor response. Participants completed simple RT (SRT), choice RT (CRT), and go/no-go (GNG) tasks, while TMS was applied at various times between the warning signal and go-signal. It was hypothesized that if MEP suppression relates to preparation level, the greatest suppression would be observed during the SRT and GNG tasks, as these paradigms encourage advance preparation and response inhibition. Conversely, if the reduction in corticospinal excitability is associated with facilitating response initiation processes, then suppression would be expected for all tasks, including the CRT paradigm in which preparation does not occur until presentation of the go-signal. Results showed MEP amplitudes decreased for all tasks as the go-signal approached; however, both the SRT and GNG had significantly greater MEP suppression 50 ms prior to, and coincident with the go-signal. These results indicate that the nature and origin of the suppression is likely multifactorial and relates to both preparatory and initiation-related processes, with the timeline and magnitude of suppression dependent on the nature of the task being executed.

**Impact Statement:** Transcranial magnetic stimulation was used to elicit motor-evoked potentials to examine the timeline of corticospinal activation during the instructed delay period for choice, simple and go/no-go reaction time tasks. For all tasks, corticospinal excitability was initially elevated compared to baseline, followed by a similar magnitude of early suppression. However, just prior to the go-signal, those tasks that allowed advance preparation showed additional suppression, providing novel information linking pre-movement corticospinal suppression to preparatory and inhibition processes.

## Introduction

Although much is still unknown about the neural planning that precedes the production of motor responses, it is clear that these preparatory processes are accompanied by changes in neural activation levels in motor-related cortical structures. For example, neural recordings from non-human primates have shown that motor cortical-related activity increases during the preparatory period prior to execution of a known action (Tanji and Evarts 1976), likely in order to facilitate fast and accurate responses. In humans, measurement of preparatory cortical activity is more challenging as direct measurement is invasive, and neuroimaging techniques often result in poor spatial and/or temporal resolution; thus, transcranial magnetic stimulation (TMS) has been used as an alternative, indirect method to quantify and examine motor preparation processes. TMS is a neurostimulation technique that uses a magnetic pulse to directly activate corticomotor neurons in the primary motor cortex (M1), which then project via the corticospinal tract to motor neurons innervating the target muscle. This descending activation can be measured using electromyography (EMG) as a motor-evoked potential (MEP). The amplitude of the MEP is thought to reflect the excitability of the corticospinal pathway with larger MEP amplitudes reflecting greater activation in the system (see Reis et al. 2008; Spampinato et al. 2023 for reviews). During a simple reaction time (RT) task where the required response is known in advance and thus can presumably be preprogrammed, it has been shown that MEP amplitudes increase from baseline levels during response preparation, and these levels are maintained until about 300 ms prior to the go signal (Kennefick et al. 2014). Interestingly, at this point, MEP amplitudes have consistently been shown to *decrease* in the short interval leading up to the go-signal (Duque et al. 2017; Greenhouse et al. 2015; Touge et al. 1998), after which they once again increase due to execution of the prepared response (Chen et al. 1998; Leocani et al. 2000; Rossini et al. 1988).

The relatively large MEP amplitudes observed during the early planning and late response initiation phases of a motor task, which mirror modulation of firing rates directly recorded in non-human primates (Churchland et al. 2006; Riehle and Requin 1989), are thought to reflect an expected increase in corticospinal activation that is associated with response preparation and execution, respectively (Davranche et al. 2007; Klein-Flügge et al. 2013). However, the reason for the observed *decrease* in MEP amplitude occurring immediately prior to the imperative signal has been a matter of debate (see Duque et al. 2017; Greenhouse 2022 for reviews). Early explanations considered pre-movement corticospinal inhibition necessary to either suppress competing movements (i.e., “competition resolution” model; Burle et al. 2004) or to maintain a high level of advance preparation while preventing the premature release of the prepared motor plan (i.e., “impulse control” model: Duque et al. 2010; Duque and Ivry 2009). However, these inhibition-related explanations have been challenged by the co-existence of decreased corticospinal excitability in task-irrelevant muscles, which would not be affected if the inhibition was strictly related to suppressing unwanted responses (Duque et al. 2017; Greenhouse et al. 2015). This observation led to an alternative “spotlight control” hypothesis whereby neural inhibition acts to speed movement initiation by increasing the signal-to-noise ratio for motor output pathways, thus facilitating response selection and execution (Greenhouse et al. 2015; Lebon et al. 2019). Further supporting an initiation-based explanation, suppression of corticospinal excitability has been shown to follow a similar time course for both reactive and self-initiated responses (Ibáñez et al. 2020). These authors argued that movements that are self-initiated without an external cue should not require any inhibitory processes related to avoiding false starts since, by definition, false starts cannot occur in this circumstance. Instead, they argued that the observed suppression may instead be a function of shifting activation from preparatory processes to response initiation processes. This explanation is also supported by the somewhat paradoxical finding that the magnitude of MEP suppression has been shown to be correlated with response latency, with *greater inhibition* associated with *shorter RTs*, suggestive of a role of preparatory inhibition in the facilitation of fast movement initiation (Hannah et al. 2018).

These previous studies have often employed one type of RT paradigm in isolation, or have not directly compared the timeline of MEP suppression between tasks; yet important information may be gleaned by comparing cortical activation dynamics between tasks that result in differences in the timeline of preparation, inhibition, and initiation processes. These types of tasks were originally considered in Donders’ “On the Speed of Mental Processes” (1969 translation of original 1868 text), with the use of simple, choice, and go/no-go RT tasks being used to calculate the durations of the various stages of information processing. In the simple RT task, the participant knows the required response in advance and can actively prepare the movement before the imperative stimulus. However, while this may enable fast RTs, this ability to prepare to a high level may necessitate greater levels of inhibition prior to the imperative stimulus to prevent premature release of the response. In contrast, when the RT task involves a response that is specified by the imperative stimulus (choice RT), the exact response is unknown in advance, which may result in a lowered state of preparation as response selection and programming is completed following the go-signal. Lastly, the go/no-go RT task involves a single known response that is either initiated in response to a “go” imperative cue or withheld in response to a “no-go” imperative cue, and as such the go/no-go task allows for response preparation to occur in advance, similar to a simple RT paradigm. However, the need to inhibit response execution on no-go trials would be expected to involve additional inhibitory processes once the imperative cue is presented, compared to simple RT in which a response is required every trial. Although the neural processes involved are undoubtedly more complex than summarized above, robust RT differences between the tasks presumably reflect (at least in part) differences in preparatory and inhibitory activation (Carlsen et al. 2004; Filipović et al. 1997), which may manifest in MEP amplitude differences throughout the RT paradigm.

The purpose of the present study was to examine the timeline of corticospinal excitability during the preparatory phase of simple, choice, and go/no-go RT tasks to provide further insight into the nature of premovement suppression of MEP amplitude. This was done by measuring TMS-evoked MEP amplitudes in the same effector muscle at various times prior to the imperative stimulus in RT tasks in which: 1) the effector muscle could be prepared and was certain to be used as a prime mover (i.e., simple RT), 2) the effector could be prepared, but this prepared action was either initiated or inhibited (i.e., go/no-go RT), and 3) the effector could not be fully prepared as the response was unknown, and may or may not need to be engaged (i.e., choice RT). If the previously observed reduction in corticospinal excitability relates to high levels of advance preparation that require inhibition to prevent premature/incorrect responses, it would be expected that simple and go/no-go RT tasks would result in larger MEP amplitude decreases prior to the imperative stimulus, as compared to the choice RT task. Conversely, if the reduction in corticospinal excitability is associated with response initiation processes, then suppression would be expected for all tasks, including the choice RT paradigm in which final preparation does not occur until presentation of the go-signal. It is also worth noting that inhibition versus initiation explanations for premovement suppression are likely not mutually exclusive and do not need to be considered as competing hypotheses, as both processes may contribute to the reduction in MEP amplitudes just prior to response execution.

## Methods

### Participants

Thirty right-handed or ambidextrous adults (17 female; mean age 26.1 years, SD = 9.6) volunteered to participate in this study. All participants had normal or corrected-to-normal vision, had no known sensory or motor dysfunctions, and were naïve to the hypotheses under investigation. The single testing session lasted approximately 90 minutes. Before the start of a testing session, participants provided written informed consent and completed a safety questionnaire to screen for contraindications to the application of TMS (Rossi et al. 2011). This study and the protocols used were approved by the University of Ottawa Research Ethics Board (File number H-05-22-8081) and was conducted in accordance with the seventh revision of the Declaration of Helsinki.

### Task & Apparatus

Participants sat approximately 1.5m in front of a 24” computer monitor (Asus VG248; 144 Hz refresh) with their right forearm secured in a custom manipulandum which restricted movement of the wrist to flexion and extension (Figure 1). Participants performed ballistic wrist extension and flexion movements within three separate RT paradigms: a simple RT task, a choice RT task, and a go/no-go RT task. Each trial began with the presentation of a fixation cross in the centre of the otherwise blank grey screen for 5 – 7 s, followed by the presentation of a visual warning signal. For the simple and go/no-go tasks the visual warning signal consisted of the appearance of single box (7 cm square, grey fill with a 3 mm black outline) presented 10 cm to the right of the fixation cross. For the choice task the visual warning signal was two identical boxes (as above) each presented 10 cm either side of the fixation cross. The warning signal was followed by a fixed foreperiod of 500 ms prior to the imperative stimulus. In the simple RT task, participants were instructed to perform a 20-degree right wrist extension presentation of the imperative go-signal which consisted of the on-screen box filling in green. In the go/no-go RT task participants were instructed to execute a 20-degree right wrist extension if a green go-signal was presented, but to withhold any motor response if the box turned red. In the choice RT task participants were instructed to perform a 20-degree wrist extension if the box on the right of the fixation cross turned green or a 20-degree wrist flexion if the box on the left of the fixation cross turned green. Participants were instructed to execute their movements as soon as possible following the presentation of the go-stimulus.

**Figure 1.**
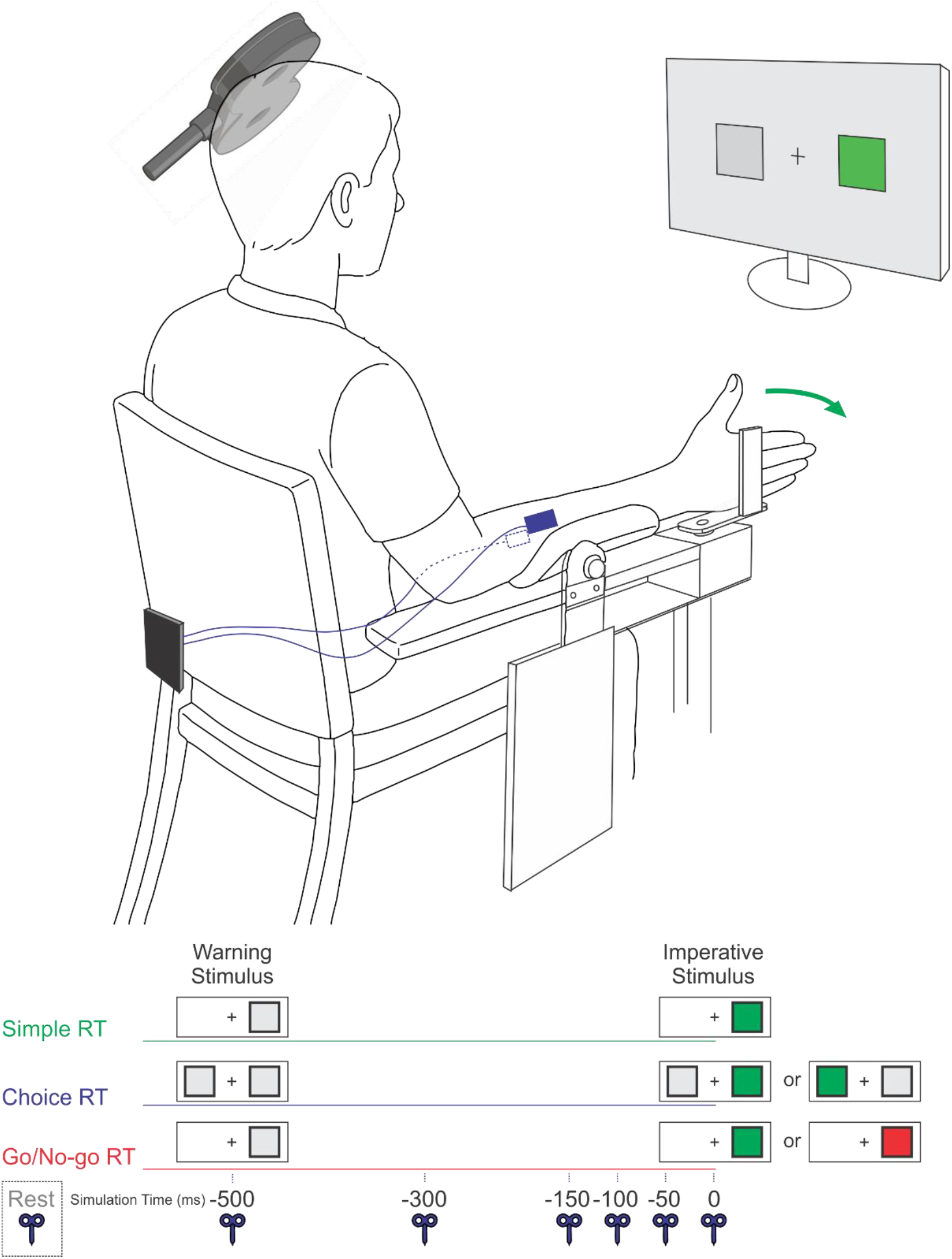
Schematic of experimental set up (top panel). Participants performed a right wrist extension (or flexion on choice trials) in response to a visual stimulus. TMS was applied over the left motor cortex and EMG data was collected from the right wrist extensors and flexors (blue). Schematic timeline, including the warning stimulus (-500 ms) and imperative stimulus (0 ms) for the various reaction time (RT) tasks, is shown at the bottom. TMS stimulation times within the time course of a trial are indicated with a TMS figure-8 coil icon.

For each RT task, participants completed a single block of 70 experimental trials; block order was counterbalanced. Prior to each RT task block, participants completed 10 familiarization trials of that task. In the simple RT task, 5 additional catch trials were randomly included in which the imperative go-stimulus did not occur. These were included in order to minimize anticipation of the go-stimulus. In the go/no-go and choice RT tasks, participants completed 35 randomized trials of each task alternative. Within each RT task block participants were given the option to rest every 25 trials to minimize fatigue.

### Transcranial Magnetic Stimulation

TMS was delivered during the reaction time tasks using a Magstim 200^2^ stimulator with a 70-mm figure-8 coil (Magstim Company Ltd., Whitland, UK) that was placed over the primary motor cortex (M1) representation of the right extensor carpi radialis (ECR) muscle. The target location was first approximated by finding the midpoint between a participant’s nasion and inion (in the midsagittal plane) and left and right pre-auricular notches (in the frontal plane). From this point, a location 4 cm lateral (leftward) and 1 cm anterior was used as a starting point to determine the TMS location over M1 that generated the largest motor-evoked potential (MEP) in the right ECR (i.e., hotspot). During stimulation, the TMS coil was oriented such that the handle pointed backwards at approximately 45-deg relative to the mid-sagittal line of the head. Neuro-navigation hardware and software (ANT Neuro Visor 2, Madison, WI) was used to save and replicate this location for all subsequent TMS applications. After the hotspot was located, the resting motor threshold (rMT) for the right ECR muscle was determined to the nearest 1% of stimulator output by finding the minimum intensity needed to elicit an MEP of 50 µV in five of ten trials (Rossini et al. 2015). During the RT tasks TMS was applied at 120% of each individual’s rMT.

On each trial (except simple RT catch trials) TMS was applied over the motor hotspot for the ECR at one of six time points with respect to the onset of the imperative go-stimulus: -500 ms (at warning stimulus onset), -300 ms, -150 ms, -100 ms -50 ms and 0 ms (coincident with the go-signal). The TMS was only applied at one of these time points per trial, and each time point was repeated 10 times (random order). For the choice RT and go/no-go tasks there were two possible response alternatives, therefore each TMS time occurred for each eventual response type on 5 separate trials. It was expected that response type (e.g., flexion vs. extension during choice RT task; “go” versus “no go” trials) should have no effect on MEP amplitude as the TMS application occurred prior to the imperative stimulus when the eventual required response was still unknown and therefore the preparatory/inhibitory processes would be expected to be similar on average at this time.

TMS was also applied on 10 trials for each task at -2500 ms (2000 ms prior to the warning stimulus) as it was assumed that MEP amplitude at this time would act as an index of “resting” baseline level activation. However, preliminary analysis indicated that there was no significant difference in MEP amplitude (p = .08) between the -2500 ms (between-trial) application time and the -500 ms application time (at the warning signal), indicating that in between trials participants remained in a task set that resulted in elevated corticospinal excitability similar to that observed at the -500 ms time point. As such, trials where TMS was applied at -2500 ms were not able to be used as resting baseline, and were discarded from analysis. To obtain a more accurate estimate of resting baseline corticospinal activation, 10 MEPs were collected in a subset of individuals (n = 12) immediately following determination of rMT with TMS output set to 120% of rMT for that individual. These MEPs were collected prior to providing any of the RT task instructions, and participants were instructed to sit quietly with their right arm resting comfortably in their lap.

### Recording Equipment

Surface electromyography (EMG) was collected from the extensor carpi radialis (ECR) and flexor carpi radialis (FCR) muscles using bipolar pre-amplified double differential surface electrodes (Delsys, Bagnoli DE-3.1; Delsys Inc., Natick, MA) connected to an external amplifier system (Delsys Bagnoli-8) via shielded cabling. The location of EMG electrodes was cleaned using abrasive gel (Nuprep) and an alcohol swap to decrease electrical impedance, with electrodes placed parallel to the muscle fibers and attached to the skin via double-sided adhesive tape. A reference electrode was placed on the medial epicondyle or olecranon of the right humerus. Raw band-passed (20-450 Hz) EMG was digitally sampled at 4 kHz (PCIe-6321, National Instruments) for 4.5 s starting 3 s prior to the imperative stimulus on each trial using a custom program written in LabVIEW software (National Instruments Inc.).

### Data Reduction

EMG and MEP parameters were determined from the raw data using a custom LabVIEW analysis program. Integrated MEP amplitude (iMEP) was calculated by numerically integrating the rectified EMG values recorded from the ECR in a 30 ms window beginning 15 ms following the time of TMS presentation. Integrated area under the curve has been argued to be a more valid amplitude measure for polyphasic MEPs which are typically seen in non-hand effectors (Spampinato et al. 2023). Premotor reaction time (RT) was defined as the time from presentation of go-stimulus to EMG onset of the agonist muscle and was only analyzed for the wrist extension task (go trials only) as it was the common movement task across all RT task blocks. EMG onset was determined as the first point at which full-wave rectified and filtered EMG data (dual-pass filtered using a 25 Hz low-pass second-order elliptic filter) increased more than 2 standard deviations above the mean value recorded during the 500 ms prior to the warning signal and remained elevated for a minimum of 20 ms. EMG onsets were visually inspected and manually adjusted if necessary.

Practice trials were not included in the analyses, nor were catch trials that occurred during the simple RT task block. Trials with the following errors were excluded from analysis: Anticipation (256 trials) was defined as trials where EMG onset of the agonist muscle occurred in less than 50 ms following a go-stimulus. Slow RT (85 trials) was defined as EMG onset of the agonist muscle occurring more than 2.5 SD greater than the mean for each RT task. Movement errors (109 trials) consisted of incorrect responses during choice and go/no-go RT tasks. Finally, trials were removed if peak-to-peak MEP amplitude was less than 50 µV (38 trials), or if there was excessive background EMG activity (RMS > 3x baseline) prior to the TMS pulse (89 trials). Therefore, the final testing trial inclusion rate for analysis was 4943/5520 (89.5%).

### Statistical Analysis

Premotor Reaction time was analyzed for the right wrist extension movement between RT tasks and TMS presentation times using a linear mixed effects model with interacting fixed factors of TMS time prior to the go-signal (6 levels: −500, −300, −150, −100, −50, 0) and RT task (3 levels: simple RT, choice RT, go/no-go RT), with random intercepts specified for participant, and random slopes specified for the effect of RT task condition (e.g., RT ∼ TMStime * Task + (1 + Task | Participant)).

To determine if RT task type affected the degree to which corticospinal excitability was elevated above the resting state, integrated MEP amplitude was compared between resting baseline (measured in subset of 12 participants; see above) and when TMS was applied at the warning signal (i.e., “go” minus 500 ms) for each of the three RT conditions. These values were compared using a generalized linear mixed effects model with task condition (4 levels: Resting Baseline, simple RT, choice RT, go/no-go RT) as a fixed factor. Random intercepts were specified for participants, and the model used a Gamma distribution for the residuals with a log link function (e.g., MEP amplitude ∼ TaskCondition + (1 | Participant), family = Gamma(link=“log”)). A Gamma model was chosen because integrated MEP amplitudes exhibited strong positive skew (Ng and Cribbie 2017).

To investigate how corticospinal excitability evolved as the go-signal approached, MEP amplitude in the wrist extensor was analyzed using a similar generalized linear mixed effects model with interacting fixed factors of TMS time prior to the go-signal (6 levels: −500, −300, −150, −100, −50, 0) and RT task (3 levels: simple RT, choice RT, go/no-go RT). Random intercepts were specified for participants, and the model used a Gamma distribution for the residuals with a log link function (e.g., MEP amplitude ∼ TMStime * Task + (1 | Participant), family = Gamma(link=“log”)).

The R software package (version 4.5.2; R Core Team 2025) was used for all analyses. The lme4 (Bates et al. 2026) lmerTest (Kuznetsova et al. 2026) packages were used to fit the models and calculate p-values. The anova() function was used to calculate p-values for main effects and interactions using Satterthwaite’s method. Type II Wald chi-squared tests were used to calculate p-values for main effects and interactions in Gamma models, The emmeans (Lenth et al. 2026) package was used for pairwise contrasts with Tukey’s HSD Post-hoc correction for multiple comparisons. The DHARMa (Hartig et al. 2024) and flexplot (Fife and A 2022) packages were used to assess assumptions of the models and to evaluate residuals. If the specified models exhibited convergence failures that could not be remedied via optimization algorithms, the models were simplified by removing random slopes. All data are expressed as estimated marginal means with standard error or 95% confidence intervals. The sjPlot (Lüdecke et al. 2025) and flexplot (Fife and A 2022) packages were used to generate visuals.

## Results

### Reaction Time

Premotor RT as a function of TMS application time (-500 ms, -300 ms, -150 ms, -100 ms, -50 ms, 0 ms) for each RT task (simple, choice, go/no-go) is presented in Figure 2. Analysis revealed a significant main effect of RT task, *F*(2, 28.7) = 55.014, p < .001), indicating that RT for the simple RT task (204 ms, SE = 6.0 ms) was significantly shorter than both go/no-go task (260 ms, SE = 5.3 ms) and the choice RT task (251 ms, SE = 6.3 ms), while there was no difference between the latter two. There was also a main effect of TMS time, *F*(5, 3059.7) = 2.544, p = .026, with post-hoc tests revealing that RT was significantly shorter (10 ms, SE = 3.4) when TMS was applied at 150 ms prior to the go versus at the go signal (p = .036). Finally, there was no significant interaction between the factors (p = .864).

**Figure 2.**
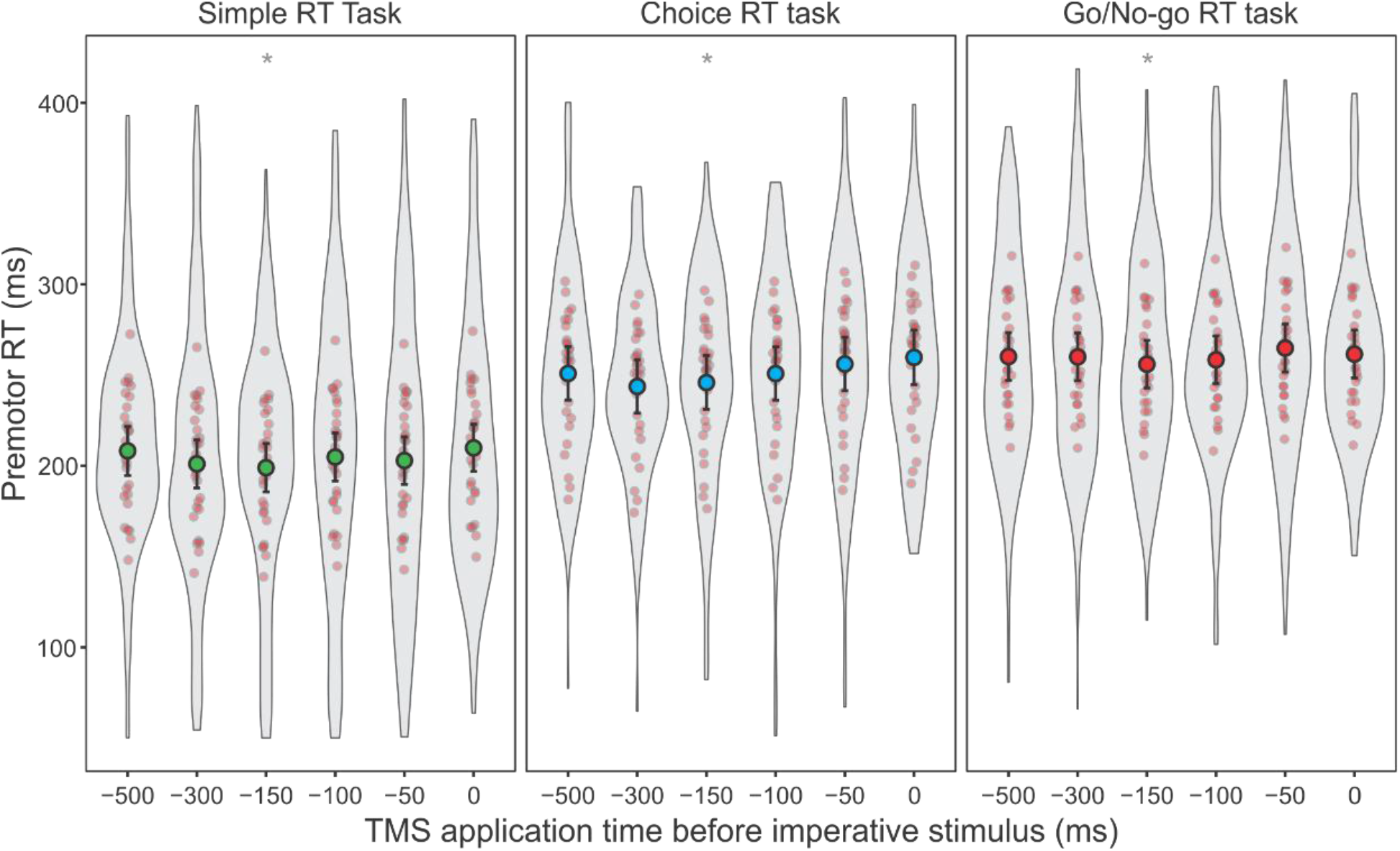
Premotor reaction time (RT) as a function of TMS stimulation time for each RT task (simple RT, choice RT, go/no-go RT). Fixed effects means and 95% CI are shown with large black markers and error bars. Individual subject estimated mean RTs are shown as small red markers, and violin plots represent the distribution of raw data, Note, RT was significantly shorter in the simple RT condition compared to choice and go/no-go. There was also a significant reduction in RT in each task when TMS was applied at the -150ms time point as compared to time zero (shown with asterisk *).

### MEP amplitude

Integrated MEP amplitude when TMS was applied at rest (prior to beginning any RT task; n=12) was compared to MEP amplitude when TMS was applied at the warning signal in each of the RT tasks (n=30). This was done to determine if corticospinal excitability was differentially elevated from baseline between the RT tasks (Figure 3). Analysis showed a significant effect of condition, χ^2^ (3) = 146.22, p < .001, with post-hoc comparisons indicating that while MEP amplitude was significantly larger in all three RT tasks compared to baseline (all p-values < .001), there was no difference in amplitude between the RT tasks (all p-values > .743).

**Figure 3.**
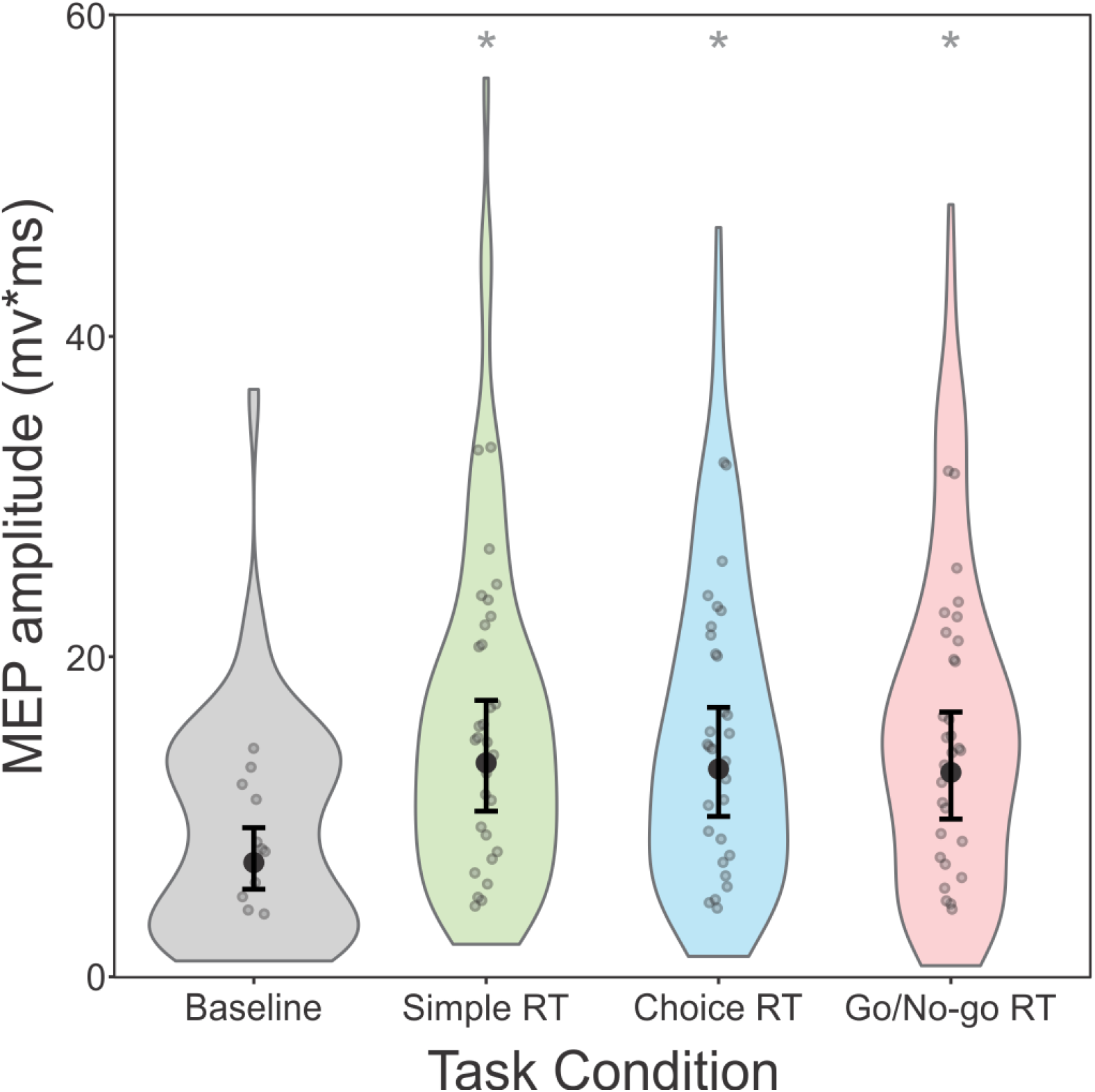
Motor-evoked potential (MEP) amplitude at baseline (resting; grey), and at the warning signal (-500 ms) for each reaction time (RT) task (green = simple RT; blue = choice RT; red = go/no-go RT). Fixed effect means and 95% CI are shown as large markers and error bars, individual subject estimated mean MEP amplitudes are shown as small grey markers, and violin plots represent the distribution of raw data, Means are back-transformed from the log scale. Note that MEP amplitude was significantly (*) larger for all three RT tasks compared to baseline.

Integrated MEP amplitude as a function of TMS time during each RT task is presented in Figure 4. Significant main effects were found for TMS time, χ^2^ (5) = 267.587, p < .001, and RT task, χ^2^ (2) = 16.980, p < .001, as well as a significant interaction between TMS time and task, χ^2^ (10) = 18.664, p = .045). Post-hoc tests indicated that in each RT task, MEP amplitude was significantly smaller (all p-values < .01) when TMS was applied at -150 ms, - 100 ms, -50 ms, and at the go-signal, as compared to when compared to when TMS was applied at -500 ms and -300 ms. Furthermore, in the Go/No-go RT task, MEP amplitude was significantly smaller at -50 ms and at the go-signal compared to -150 ms (p-values < .002).

**Figure 4.**
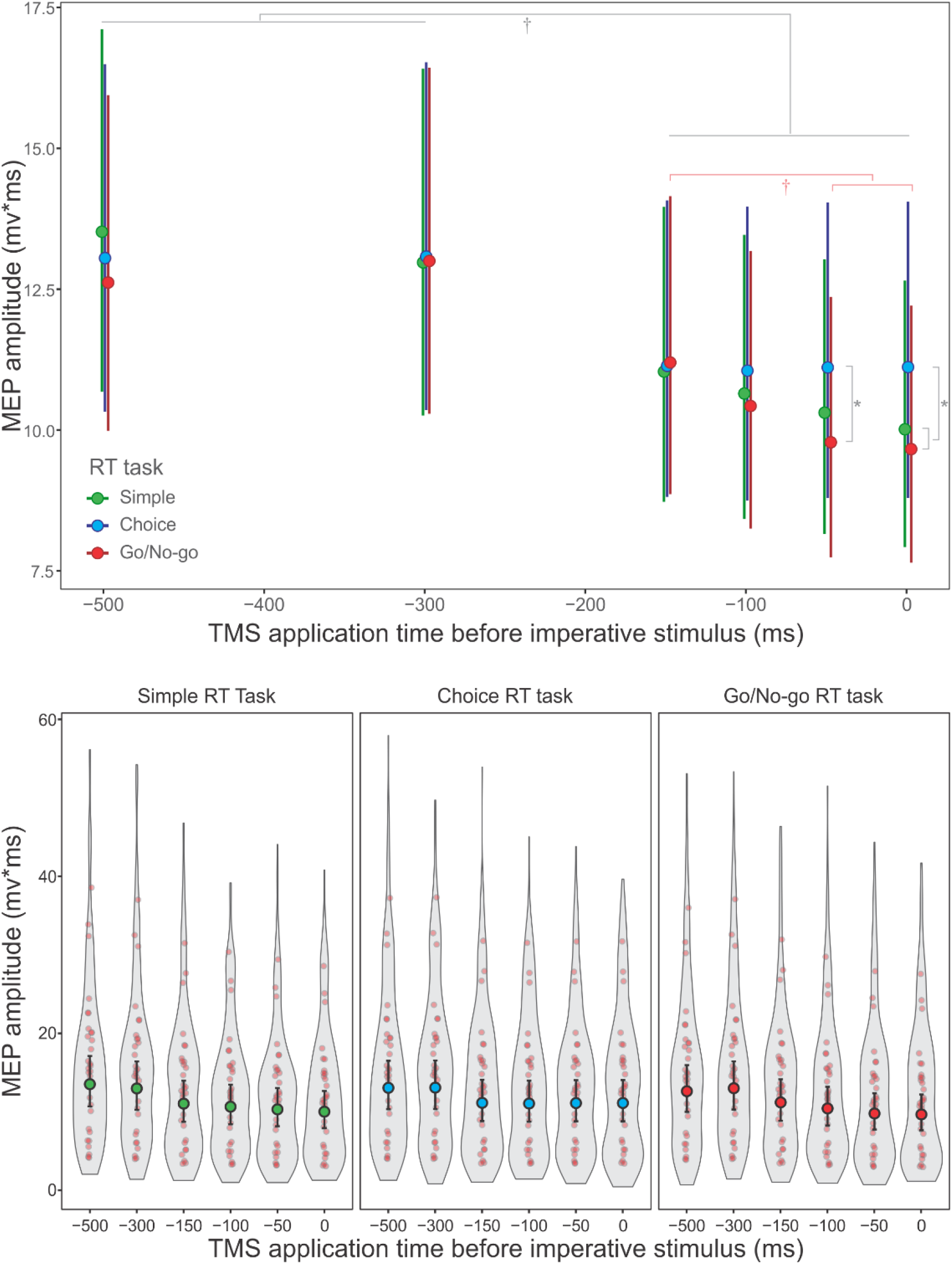
Motor-evoked potential (MEP) amplitude as a function of TMS stimulation time for each reaction time (RT) task (green = simple RT; blue = choice RT; red = go/no-go RT). Top panel shows fixed effect means and 95% CI for each task overlayed and scaled with time for ease of comparison. Bottom panel shows fixed effects means and 95% CI with large markers and error bars in separate panels for each task. Individual subject estimated mean MEP amplitudes are shown as small red markers, and violin plots represent the distribution of raw data. Note that means are back-transformed from the log scale. Asterisk (*) indicates significant difference between tasks at each time point. Dagger (†) indicates significant difference between time points (grey = all tasks, red = go/no only).

When comparing between RT tasks at each TMS time point, MEP amplitude for the choice RT task was significantly larger than that for the go/no-go RT task (p < .001) when TMS was applied either 50 ms before or coincident with the go-signal. Similarly, MEP amplitude for the choice RT task was significantly larger than that for the simple RT task when TMS was applied coincident with the go-signal (p = .010), but did not reach significance for the 50 ms time point (p = .091).

## Discussion

The present study investigated how the time course of premovement suppression of corticospinal excitability (CE) was affected by the type of reaction time (RT) task being performed (simple RT, choice RT, go/no-go RT). All three RT tasks required the participant to perform a similar wrist extension movement on some trials, but under different cueing conditions. Because the level of certainty regarding the required response differs between these tasks, it has been argued that differences exist in the level of advance preparation that is afforded by each task (Leuthold et al. 2004), as well as the required level of inhibitory activation to restrain unwanted movement (Greenhouse et al. 2015). These differences appeared to be reflected in the RT results, as shown by significantly shorter response latency for the simple RT task (Figure 2), as this condition allowed for 1) more advance preparation than choice RT, and 2) no need to inhibit responses on “no go” trials. Most importantly, the type of RT task also resulted in differences in the timeline of CE suppression during the delay period. As compared to rest, corticospinal excitability was similarly elevated at 500 ms prior to the go-signal in all tasks, indicating the presence of general preparatory task-related neural activity (Figure 3). Although suppression of this excitability was observed in all three RT tasks beginning at approximately 150 ms prior to the go-signal, for the simple RT and go/no-go RT tasks, CE was further suppressed as the go-signal approached, while no further suppression was observed in the choice RT task (Figure 4). These results indicate that RT task requirements, including the ability to prepare a response in advance and the potential requirement to inhibit a response, affect the timeline and magnitude of premovement suppression observed in the delay period leading up to the go-signal in different RT tasks.

### The Effect of Task Requirements and TMS on Reaction Time

As has been demonstrated for over a century (Donders 1969), the different RT tasks used here exhibited differences in mean RT with simple RT being significantly shorter than choice RT and go/no-go RT (Figure 2). The shorter RT latencies observed in simple RT tasks are typically attributed to fact that the required response is certain, and thus no response selection need occur during the RT interval. In this case, given sufficient time, the response can be fully prepared in advance, and the only requirement for the participant is to detect that the imperative stimulus has occurred and to initiate the prepared action (Niemi and Näätänen 1981). On the other hand, choice RT incurs additional processing time, ostensibly due to the requirement for response selection and response preparation prior to initiation. For a go/no-go RT task the response on go-trials is certain, but additional processing must be undertaken during the RT interval to discriminate between stimuli in order to determine whether to initiate or withhold the planned response. These additional processing steps lead to longer go/no-go RT latencies compared to simple RT, as seen here. Although choice RT tasks have often been reported to exhibit longer RT than go/no-go tasks (Brebner and Welford 1980), this was not the case in the present experiment (Figure 2), and previous studies have also reported similar negligible RT differences in RT between these tasks in young adults (Gomez et al. 2007; Perea et al. 2016).

As a secondary RT finding, the application of TMS also had a small (∼10 ms) but consistent effect of reducing RT when TMS was applied at 150 ms prior to the go-signal. Similar RT speeding on TMS trials has been reported previously (Leocani et al. 2000; Pascual-Leone et al. 1992) and because this effect was significant only at 150 ms prior to the go-signal (and to a lesser extent at 300 ms prior to the go, p = .100), it likely reflects a cueing effect whereby the physical sensation associated with the TMS predicted the upcoming go-signal. This effect was not different between RT tasks, and at later stimulation times the sound / tactile stimulation from the TMS would likely have occurred too late for it to be processed as a cue.

### The Effect of Task Requirements on Corticospinal Suppression

Given the RT latency differences between conditions, it was predicted that the task-specific timeline of preparation, inhibition, and initiation processes would be reflected in the magnitude of MEP changes. The general level of corticospinal activity (as indexed by MEP amplitude) was elevated at the warning signal, as compared to baseline for all three RT tasks (Figure 3), suggesting that this increased excitability is non-specific and not strictly related to the level of preparation for a specific response. If this activity was directly related to the level of response specific preparation, then excitability would be expected to be highest for the simple RT task (where the required response is certain) and lowest for the choice RT task (where a lack of advance knowledge precludes response-specific preparation). However, it should be noted that partial advance preparation can occur in a choice RT task, depending on the specific task requirements, as well as the number and nature of the potential response alternatives (Cisek 2006). Here, response alternatives included flexion or extension with the right wrist and as such, it is possible that at least partial preparation specific to the involved limb was able to be undertaken, as compared to a “true” choice in which no aspect of the required response is known in advance (e.g., Rosenbaum 1980).

The increase in activation from the baseline condition was maintained until the -300 ms time point, at which time CE suppression occurred in all conditions, as shown by a significant decrease in MEP amplitude at the -150 ms time point (∼11-18%; Figure 4). Although this suppression prior to the go-signal replicates many previous studies (e.g., Duque et al. 2017; Greenhouse et al. 2015; Touge et al. 1998), the lack of difference between tasks during this timeframe, even though RTs differed greatly, suggests that this “early” (-300 ms to -150 ms) suppression is not due to specific preparatory or inhibitory processes associated with the upcoming response. Instead, it is likely related to the anticipation of initiation processes following the go-signal, such as increasing the signal-to-noise ratio for motor output pathways (i.e., “spotlight control”; Greenhouse et al. 2015). This assertion is also consistent with studies that concluded CE suppression relates to initiation process, due to its presence in task irrelevant muscles (Quoilin et al. 2019), as well as during self-initiated responses (Ibáñez et al. 2020).

Although the initial CE suppression was not task dependent, in the final 150 ms prior to the go-signal, differences occurred between the RT tasks (Figure 4). Specifically, the conditions that allowed for advance response preparation (simple and go/no-go RT tasks) showed further reductions in MEP amplitude (9-14%;) as compared to the choice RT condition (in which the largest additional suppression was 0.8%, p > .999). This “late” suppression (-150 ms to 0 ms) does not seem to relate to the facilitation of response initiation processes, as implied by the spotlight hypothesis (Greenhouse et al. 2015) because an increased signal-to-noise ratio would potentially be *least* necessary when the response is known in advance. While the simplest explanation might be that this additional suppression in the simple and go/no-go tasks is associated with inhibitory processes that serve to prevent the prepared response from being released prior to the go-signal (e.g., “impulse control”; Duque et al. 2010; Duque and Ivry 2009), an impulse control explanation cannot fully account for some previous findings. For example, Ibáñez et al. (2020) demonstrated that suppression was observed even in self-initiated responses, where impulse control would be unnecessary. Instead, the late suppression observed here in the simple RT and go/no-go tasks may be more consistent with the explanation that the suppression reflects a system that is closer to a state that is ready for a response to be executed (Hannah et al. 2018; Ibáñez et al. 2020). That is, both the simple RT task and the go/no-go task enabled a greater state of preparation of the specific motor task (if needed), leading to larger observed suppression.

It may seem unexpected that a similar level of CE suppression was seen between the simple RT and go/no-go tasks, as the RTs were very different (with significantly longer RT latency in the go/no-go conditions; Figure 3). However, the RT difference is not surprising because although both tasks likely involve advance preparation and some inhibitory processes to prevent false starts, the go/no-go task likely requires additional processes once the imperative stimulus occurs. Specifically, the individual must determine whether a “go” signal occurred or a “no-go” signal occurred (and thus release, maintain, or increase any response-related inhibitory processes), which may not be reflected in the delay interval CE activation level. Importantly, this RT difference also suggests that the suppression seen in these two conditions may originate from different mechanisms: it is possible that in the go/no-go task, a greater proportion of the observed suppression was related to “impulse control” whereby suppression acts to prevent unwanted premature or incorrect release of the prepared motor response (Duque and Ivry 2009), whereas in the simple RT task a greater amount of the suppression was related to an increased preparatory state (Ibáñez et al. 2020). Nevertheless, further study would be needed to determine the relative contributions of each of these mechanisms to the observed suppression.

### Summary

In conclusion, the present experiment utilized different reaction time tasks, which involved distinct preparation and inhibition processing requirements to examine the timing and magnitude of suppression of corticospinal excitability just prior to the go-signal. No differences were evident in the MEP amplitude at the warning signal, which then showed similar suppression for all three RT tasks until 150 ms prior to the go-signal. During the final 150 ms, the magnitude of suppression diverged between the tasks, as the choice RT resulted in no further suppression, compared to the simple and go/no-go RT tasks, which exhibited decreased MEP amplitude until the go-signal. Collectively, these results indicate that the nature and origin of the suppression is likely multifactorial, with early (300 to 150 ms prior to the go-signal) corticospinal excitability suppression potentially functioning to improve signal detection/initiation processes, and later (within 150 ms of the go-signal) suppression relating to preparatory state optimization and/or inhibitory processes to prevent early or unwanted responses. Thus, changes in corticospinal excitability may be an inherent feature of processes associated with both motor preparation and initiation, and the timeline and magnitude of suppression dependent on the nature of the task being executed.

## Data and code availability statement

Summary data supporting the findings presented here are available at the following DOI: 10.7910/DVN/QN0N2G (Harvard Dataverse; https://doi.org/10.7910/DVN/QN0N2G)

## Ethics approval statement

This study and the protocols used were approved by the University of Ottawa Research Ethics Board (File number H-05-22-8081) and was conducted in accordance with the seventh revision of the Declaration of Helsinki.

## CRediT author statement

**Anthony N. Carlsen**: Conceptualization, Methodology, Software, Formal analysis, Resources, Writing – Original draft, Writing-Reviewing and Editing, Visualization, Supervision, Project Administration, Funding acquisition. **Cassandra M. Santangelo**:: Investigation, Data curation, Writing – Original draft, Writing-Reviewing and Editing. **Christin M. Sadler**: Investigation, Data curation, Writing – Original draft, Writing-Reviewing and Editing. **Dana Maslovat**: Conceptualization, Methodology, Formal analysis, Writing – Original draft, Writing-Reviewing and Editing.

## References

Bates D, Maechler M, Bolker [cre B, aut Walker S, Christensen RHB, Singmann H, Dai B, Scheipl F, Grothendieck G, Green P, Fox J, Bauer A, simulate.formula) PNK (shared copyright on, Tanaka E, Jagan M, Boylan RD, Ly A. lme4: Linear Mixed-Effects Models using “Eigen” and S4 [Online]. 2026. https://cloud.r-project.org/web/packages/lme4/index.html [20 Apr. 2026].

Brebner JMT, Welford AT. Introduction: an historical background sketch. In: Reaction times, edited by Welford AT. London: Academic Press, 1980, p. 1–23.

Burle B, Vidal F, Tandonnet C, Hasbroucq T. Physiological evidence for response inhibition in choice reaction time tasks. Brain and Cognition 56: 153–164, 2004.

Carlsen AN, Chua R, Inglis JT, Sanderson DJ, Franks IM. Can prepared responses be stored subcortically? Exp Brain Res 159: 301–309, 2004.

Chen R, Yaseen Z, Cohen LG, Hallett M. Time course of corticospinal excitability in reaction time and self-paced movements. Ann Neurol 44: 317–325, 1998.

Churchland MM, Yu BM, Ryu SI, Santhanam G, Shenoy KV. Neural variability in premotor cortex provides a signature of motor preparation. J Neurosci 26: 3697–712, 2006.

Cisek P. Integrated neural processes for defining potential actions and deciding between them: a computational model. J Neurosci 26: 9761–9770, 2006.

Davranche K, Tandonnet C, Burle B, Meynier C, Vidal F, Hasbroucq T. The dual nature of time preparation: neural activation and suppression revealed by transcranial magnetic stimulation of the motor cortex. Eur J Neurosci 25: 3766–74, 2007.

Donders FC. On Speed of Mental Processes. Acta Psychol (Amst) 30: 412–431, 1969.

Duque J, Greenhouse I, Labruna L, Ivry RB. Physiological markers of motor inhibition during human behavior. Trends Neurosci 40: 219–236, 2017.

Duque J, Ivry RB. Role of corticospinal suppression during motor preparation. Cereb Cortex 19: 2013–24, 2009.

Duque J, Lew D, Mazzocchio R, Olivier E, Ivry RB. Evidence for two concurrent inhibitory mechanisms during response preparation. J Neurosci 30: 3793–802, 2010.

Fife, A D. Flexplot: Graphically-based data analysis. Psychological Methods 27: 19, 2022.

Filipović SR, Čovičković-Šternić N, Radović VM, Dragašević N, Stojanović-Svetel M, Kostić VS. Correlation between Bereitschaftspotential and reaction time measurements in patients with Parkinson’s disease measuring the impaired supplementary motor area function? Journal of the Neurological Sciences 147: 177–183, 1997.

Gomez P, Ratcliff R, Perea M. A Model of the Go/No-Go Task. J Exp Psychol Gen 136: 389–413, 2007.

Greenhouse I. Inhibition for gain modulation in the motor system. Exp Brain Res 240: 1295–1302, 2022.

Greenhouse I, Sias A, Labruna L, Ivry RB. Nonspecific inhibition of the motor system during response preparation. J Neurosci 35: 10675–10684, 2015.

Hannah R, Cavanagh SE, Tremblay S, Simeoni S, Rothwell JC. Selective Suppression of Local Interneuron Circuits in Human Motor Cortex Contributes to Movement Preparation. J Neurosci 38: 1264–1276, 2018.

Hartig F, Lohse L, leite M de S. DHARMa: Residual Diagnostics for Hierarchical (Multi-Level / Mixed) Regression Models [Online]. 2024. https://cloud.r-project.org/web/packages/DHARMa/index.html [20 Apr. 2026].

Ibáñez J, Hannah R, Rocchi L, Rothwell JC. Premovement suppression of corticospinal excitability may be a necessary part of movement preparation. Cerebral Cortex 30: 2910–2923, 2020.

Kennefick M, Maslovat D, Carlsen AN. The time course of corticospinal excitability during a simple reaction time task. PLoS ONE 9: e113563, 2014.

Klein-Flügge MC, Nobbs D, Pitcher JB, Bestmann S. Variability of Human Corticospinal Excitability Tracks the State of Action Preparation. J Neurosci 33: 5564–5572, 2013.

Kuznetsova A, Brockhoff PB, Christensen RHB, Jensen SP. lmerTest: Tests in Linear Mixed Effects Models [Online]. 2026. https://cloud.r-project.org/web/packages/lmerTest/index.html [20 Apr. 2026].

Lebon F, Ruffino C, Greenhouse I, Labruna L, Ivry RB, Papaxanthis C. The Neural Specificity of Movement Preparation During Actual and Imagined Movements. Cereb Cortex 29: 689–700, 2019.

Lenth RV, Piaskowski J, Banfai B, Bolker B, Buerkner P, Giné-Vázquez I, Hervé M, Jung M, Love J, Miguez F, Riebl H, Singmann H. emmeans: Estimated Marginal Means, aka Least-Squares Means [Online]. 2026. https://cloud.r-project.org/web/packages/emmeans/index.html [20 Apr. 2026].

Leocani L, Cohen LG, Wassermann EM, Ikoma K, Hallett M. Human corticospinal excitability evaluated with transcranial magnetic stimulation during different reaction time paradigms. Brain 123: 1161–1173, 2000.

Leuthold H, Sommer W, Ulrich R. Preparing for action: Inferences from CNV and LRP. J Psychophysiol 18: 77–88, 2004.

Lüdecke D, Bartel A, Schwemmer C, Powell C, Djalovski A, Titz J. sjPlot: Data Visualization for Statistics in Social Science [Online]. 2025. https://cloud.r-project.org/web/packages/sjPlot/index.html [20 Apr. 2026].

Ng VKY, Cribbie RA. Using the Gamma Generalized Linear Model for Modeling Continuous, Skewed and Heteroscedastic Outcomes in Psychology. Curr Psychol 36: 225–235, 2017.

Niemi P, Näätänen R. Foreperiod and simple reaction-time. Psychol Bull 89: 133–162, 1981.

Pascual-Leone A, Valls-Solé J, Wassermann EM, Brasil-Neto J, Cohen LG, Hallett M. Effects of focal transcranial magnetic stimulation on simple reaction time to acoustic, visual and somatosensory stimuli. Brain 115: 1045–59, 1992.

Perea M, Devis E, Marcet A, Gomez P. Are go/no-go tasks preferable to two-choice tasks in response time experiments with older adults? Journal of Cognitive Psychology 28: 147–158, 2016.

Quoilin C, Fievez F, Duque J. Preparatory inhibition: Impact of choice in reaction time tasks. Neuropsychologia 129: 212–222, 2019.

R Core Team. R: A Language and Environment for Statistical Computing [Online]. R Foundation for Statistical Computing. https://www.R-project.org/.

Reis J, Swayne OB, Vandermeeren Y, Camus M, Dimyan MA, Harris-Love M, Perez MA, Ragert P, Rothwell JC, Cohen LG. Contribution of transcranial magnetic stimulation to the understanding of cortical mechanisms involved in motor control. The Journal of Physiology 586: 325–51, 2008.

Riehle A, Requin J. Monkey primary motor and premotor cortex: single-cell activity related to prior information about direction and extent of an intended movement. Journal of Neurophysiology 61: 534–549, 1989.

Rosenbaum DA. Human movement initiation: specification of arm, direction, and extent. J Exp Psychol Gen 109: 444–474, 1980.

Rossi S, Hallett M, Rossini PM, Pascual-Leone A. Screening questionnaire before TMS: An update. Clinical Neurophysiology 122: 1686, 2011.

Rossini PM, Burke D, Chen R, Cohen LG, Daskalakis Z, Di Iorio R, Di Lazzaro V, Ferreri F, Fitzgerald PB, George MS, Hallett M, Lefaucheur JP, Langguth B, Matsumoto H, Miniussi C, Nitsche MA, Pascual-Leone A, Paulus W, Rossi S, Rothwell JC, Siebner HR, Ugawa Y, Walsh V, Ziemann U. Non-invasive electrical and magnetic stimulation of the brain, spinal cord, roots and peripheral nerves: Basic principles and procedures for routine clinical and research application. An updated report from an I.F.C.N. Committee. Clinical Neurophysiology 126: 1071–1107, 2015.

Rossini PM, Zarola F, Stalberg E, Caramia M. Pre-movement facilitation of motor-evoked potentials in man during transcranial stimulation of the central motor pathways. Brain Res 458: 20–30, 1988.

Spampinato DA, Ibanez J, Rocchi L, Rothwell J. Motor potentials evoked by transcranial magnetic stimulation: interpreting a simple measure of a complex system. The Journal of Physiology 601: 2827–2851, 2023.

Tanji J, Evarts EV. Anticipatory activity of motor cortex neurons in relation to direction of an intended movement. J Neurophysiol 39: 1062–8, 1976.

Touge T, Taylor JL, Rothwell JC. Reduced excitability of the cortico-spinal system during the warning period of a reaction time task. Electroencephalogr Clin Neurophysiol 109: 489–495, 1998.

